# Utility of alternate stable isotope calibration equations and informative priors when estimating the breeding origin of Arctic-breeding shorebirds

**DOI:** 10.1101/2025.10.25.684473

**Authors:** Devin R. de Zwaan, Julie Paquet, Rebeca C. Linhart, Erica Nol, Paul A. Smith, Diana J. Hamilton

**Affiliations:** Department of Biology, Mount Allison University, Sackville, New Brunswick, Canada; Wildlife Research Division, Environment & Climate Change Canada, Delta, British Columbia, Canada; Canadian Wildlife Service, Environment & Climate Change Canada, Sackville, New Brunswick, Canada; Department of Natural Resources Science, University of Rhode Island, Kingston, Rhode Island, USA; Department of Biology, Trent University, Peterborough, Ontario, Canada; Wildlife Research Division, Environment and Climate Change Canada, Ottawa, Ontario, Canada

**Author notes:** Corresponding author: Devin R. de Zwaan.

**Keywords:** Bayesian, deuterium, eBird occupancy probability, feather isoscape, migratory connectivity, staging grounds, waders

## Abstract

Understanding migratory connectivity is essential for monitoring and conserving Arctic-breeding shorebirds, particularly given divergent rates of decline across populations. Stable hydrogen isotope analysis of feathers offers a scalable, non-invasive method to assign breeding origin, but current applications are limited by the absence of shorebird-specific calibration equations, which associate feather isotope values with environmental isotope gradients (“isoscapes”), and by longitudinal isoscape bands in the Arctic that result in diffuse origin estimates across broad, heterogeneous regions. We evaluated the utility of alternate calibration equations and the incorporation of occupancy-based informative priors for improving breeding origin assignments. Specifically, we analyzed feather isotope data from hatch-year semipalmated sandpipers (*Calidris pusilla*) captured at staging sites in the Canadian Maritimes, as well as known-origin Arctic samples, to compare calibration equations from different reference datasets and test priors derived from eBird habitat occupancy predictions. We demonstrate that Arctic samples of known origin largely align with calibration equations based on broad passerine datasets, although incorporating Arctic samples into the calibration may correct for a marginal westward bias in origin estimates. Using occupancy-based priors that reflect habitat preferences increased precision by 18.8%, identifying ‘hotspots’ within otherwise expansive isotopic bands. We highlight the validity of existing reference datasets with broad geographic coverage and demonstrate the value of informative priors to refine origin estimates. We recommend expanding Arctic sample collections across a wider longitudinal gradient to define a shorebird-specific calibration equation and further exploring potential biases in prior data sources to improve accuracy of Arctic-breeding shorebird origin assignments.

## Introduction

Arctic-breeding shorebird populations are in rapid decline globally (Birdlife International 2024). In the Americas, these declines appear spatially segregated across the breeding range, with larger declines in eastern compared to western Arctic breeding populations (Smith et al. 2023). Heterogeneity in population declines underlines the importance of accurately assigning individual shorebirds captured at southern staging and non-breeding sites to their respective Arctic breeding origins. Such knowledge allows us to quantify the relative use of staging or stopover habitats for different subpopulations, the strength of migratory connectivity and their importance for conservation, as well as changes in relative abundance and use over time (Murray et al. 2017, Morrick et al. 2022, Knight et al. 2024). While technological advances have resulted in an exponential increase in fine-scale tracking data over the past two decades, there are significant gaps in geographic, species, and population-level coverage that impact relevance for conservation across broad scales (Scarpignato et al. 2023). Stable isotope analysis allows for a more scalable, cost-effective, and non-invasive approach to establish breeding origins for a larger proportion of bird populations across more diverse regions (Hobson 2011).

Deuterium (δ^2^H) is a stable isotope that has been widely used to estimate breeding origin from measurable isotopic signatures in body tissues, like feathers (Hobson and Wassenaar 2008). This process involves associating feather deuterium levels with a deuterium ‘isoscape’, representing environmental gradients in precipitation, to spatially estimate where feathers were grown (West et al. 2010, Hobson 2011). Estimating a feather isoscape (δ^2^H _f)_ from an environmental isoscape (δ^2^H _p)_ requires a calibration equation to account for isotopic fractionation across tissues and trophic levels, which is developed with tissue samples of known origin (Hobson and Wassenaar 2008). To date, there is no calibration equation for Arctic-breeding shorebirds, and most attempts at identifying breeding origin for shorebirds use calibrations developed from passerine or waterfowl systems with known-origin samples from lower-to-mid latitudes, resulting in considerable uncertainty around assignments (Reed et al. 2018). The few available high-latitude samples of known origin belong to raptor species (Lott and Smith 2006), which, like migratory songbirds and waterfowl, represent a different trophic level to shorebirds and thus conversion of precipitation-to-feather isotope values may not be representative (Hobson 2011, Hobson et al. 2012). There is a clear need for developing shorebird-specific calibrations or evaluating the representativeness of existing equations for shorebird origin assignments.

Beyond the use of appropriate calibrations, origin assignments for species that breed across wide longitudinal ranges, like Arctic-breeding shorebirds, can be rather coarse. Several approaches have been used to refine estimates using complementary data, including multiple isotopes (Royle and Rubenstein 2004), genetic markers (Chabot et al. 2012), tracking data (Hobson and Kardynal 2016), and abundance data (Norris et al. 2006, González-Prieto et al. 2011, Hallworth et al. 2013, Fournier et al. 2017, Reese et al. 2019). These examples demonstrate the potential of probabilistic models that make use of prior knowledge (Hobson et al. 2014). Compared to abundance data, occupancy probability tends to be less variable and is particularly promising because heterogeneous landscapes within coarse regions identified by single-isotope approaches allow breeding habitat preferences to inform predictions. Specifically, a randomly sampled individual within a broad region has a greater chance of originating from areas with a higher probability of occupancy (Royle and Rubenstein 2004). Despite this, occupancy probability surfaces based on wide-scale observational data and habitat associations are relatively under-used as priors in stable isotope modelling approaches (Reese et al. 2019), particularly in a Bayesian framework that explicitly models uncertainty; a critical point given that non-Bayesian priors can propagate error (Rushing et al. 2017, Hobson 2019). Arctic shorebird breeding origins have not yet been estimated using occupancy data as informative priors due to concerns about potentially introduced bias (Reed et al. 2018), necessitating an evaluation of the utility and reliability of this approach.

We assessed the effect of different calibration equations and using occupancy probability as an informative prior when estimating breeding origin for an Arctic-breeding shorebird from feather deuterium signatures. Hatch-year semipalmated sandpiper (*Calidris pusilla*) captured during southward migration at two staging sites in Atlantic Canada were used as a case study because we can be sure feathers were grown at the breeding site. Our objectives were to: (1) compare breeding origin estimates between models with uninformative versus informative priors using occupancy probability surfaces produced from citizen science observations (i.e., eBird Status and Trends products; Fink et al. 2024); (2) compare predictions using calibration equations derived from different reference datasets of known-origin samples; (3) assess how known-origin feather samples we collected from Arctic-breeding shorebirds influenced calibration and breeding origin estimates; and (4) evaluate how our predictions aligned with known morphological gradients in semipalmated sandpipers across their breeding range as a form of ground-truthing. We expected the use of informative priors to improve precision and for known-origin Arctic-breeding shorebird samples to align more strongly with ground-foraging insectivore reference datasets in the formulation of a calibration equation. The outcomes of this study will improve future Arctic-breeding shorebird origin assignment practices and highlight next steps for future research.

## Methods

### Field methodology

We captured 143 hatch-year semipalmated sandpiper at two staging sites in New Brunswick ∼50 km apart near Petit-Cap, New Brunswick (46.193° N, -64.162° W) and Johnson’s Mills (45.834° N, -64.513° W) during southbound migration in 2018 (n = 53) and 2019 (n = 90). All captures took place between August 7 and September 13. At Johnson’s Mills we captured birds using a Fundy pull trap (Hicklin et al. 1989) and at Petit-Cap we deployed arrays of mist-nets (18 mm mesh) in the intertidal zone, perpendicular to the shore, and captured birds after dusk on a rising tide both passively and using call playbacks to attract flocks to the area. At capture, we measured morphometrics (flattened wing length, tarsus, culmen), mass, and recorded age based on plumage and feather wear following Pyle (2008). We also collected a drop of blood on a Whatman FTA card (Whatman®, Marlborough MA) through brachial venipuncture using a 27-gauge needle. Each bird was then sexed following a molecular sexing procedures adapted from Fridolfsson and Ellegren (1999). To measure deuterium stable isotopes, we collected the 6^th^ primary covert from each wing. Hatch-year plumage is completed within the first 22 days post-hatch, or before the end of July, ensuring feathers are grown on the breeding grounds (Hicklin and Gratto-Trevor 2020). A field-readable plastic flag with a three-digit alphanumeric code was affixed to the upper right leg and a metal USGS band to the upper left leg of each bird for future identification. Birds were reaccustomed to low-light conditions in a holding pen for 10 minutes before release.

In July 2022, natal down was sampled from recently fledged shorebirds of known breeding origin at three Arctic sites: Churchill, Manitoba (58.732° N, -93.797° W), East Bay (Qaqsauqtuuq) Migratory Bird Sanctuary (63.984° N, -81.707° W), and Prince Charles Island, Nunavut (68.181° N, -76.672° W). While semipalmated sandpiper feathers could not be collected at these sites, four species breeding in similar habitats were sampled: whimbrel (*Numenius hudsonicus*; n = 2), ruddy turnstone (*Arenaria interpres*; n = 6), white-rumped sandpiper (*Calidris fuscicollis*; n = 5), and semipalmated plover (*Charadrius semipalmatus;* n = 2). Since these species share similar breeding habitats and food sources, we assume their feather δ^2^H ratios should be representative of semipalmated sandpiper. Likewise, given that they are also considered income breeders (Klassen et al. 2001, Hobson and Jehl Jr 2010), we are confident that the natal down of hatch-year birds reflect the isotopic value of food intake on the breeding grounds where feathers are first grown, with little-to-no influence of imported resources through egg transfer.

### Stable isotope analysis

To prepare feathers for δ^2^H stable isotope measurement, each feather was placed in a 20 mL scintillation vial and soaked in a 2:1 Chloroform:Methanol solution for 24 hours to remove surface oils. Feathers were then rinsed twice with 2:1 Chloroform:Methanol and air dried for 48 hrs. Feather barbs were clipped from the rachis and weighed in a tared silver capsule using the Mettler-Toledo MX5 microbalance (± 0.001 mg), targeting a sample weight of 0.350 mg.

Isotope analysis was run by the Environmental Analytics and Stable Isotope Laboratory (EASIL) in Sackville, New Brunswick. Feather subsamples were combusted through flash pyrolysis at 1400°C using an Elemental Analyzer, which measured the subsamples by isotope-ratio mass spectrometry. The non-exchangeable δ^2^H was determined by comparative equilibration according to Wassenaar and Hobson (2003), based on two calibrated keratin δ^2^H isotope reference materials. All results are presented as delta values, expressed in per mille (‰), and are relative to the Vienna Standard Mean Ocean Water – Standard Light Antarctic Precipitation (VSMOW-SLAP) standard scale. We used the following calculation:

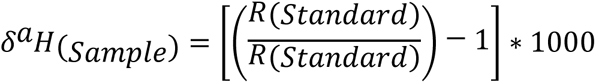

where *a* = the heavier isotope and *R* = the ratio of heavy to light isotope.

### Statistical methods

We used a single isotope (deuterium; δ^2^H) approach to identify probable natal origin of hatch-year semipalmated sandpipers captured at their staging ground during southward migration (Hobson et al. 2012). Specifically, we used a Bayesian inversion method, which fits a stochastic calibration model leveraging variance among feather deuterium samples at known locations relative to variance in an underlying environmental isoscape to produce individual probability surfaces of breeding origin (Wunder 2010). We fit an ordinary least squares linear calibration equation between an average growing-season precipitation deuterium (δ^2^H_p)_ probability surface (Bowen et al. 2005) and deuterium isotope values from feather tissue of known origin representing species breeding in North America (Hobson et al. 2012) to produce a calibrated feather tissue isoscape (δ^2^H_f;_ Figure 1). The taxa used for this calibration depended on the comparison being made (see *Testing alternative calibrations*). All known-origin feather samples were scaled to the VSMOW-H international reference scale.

**Figure 1.**
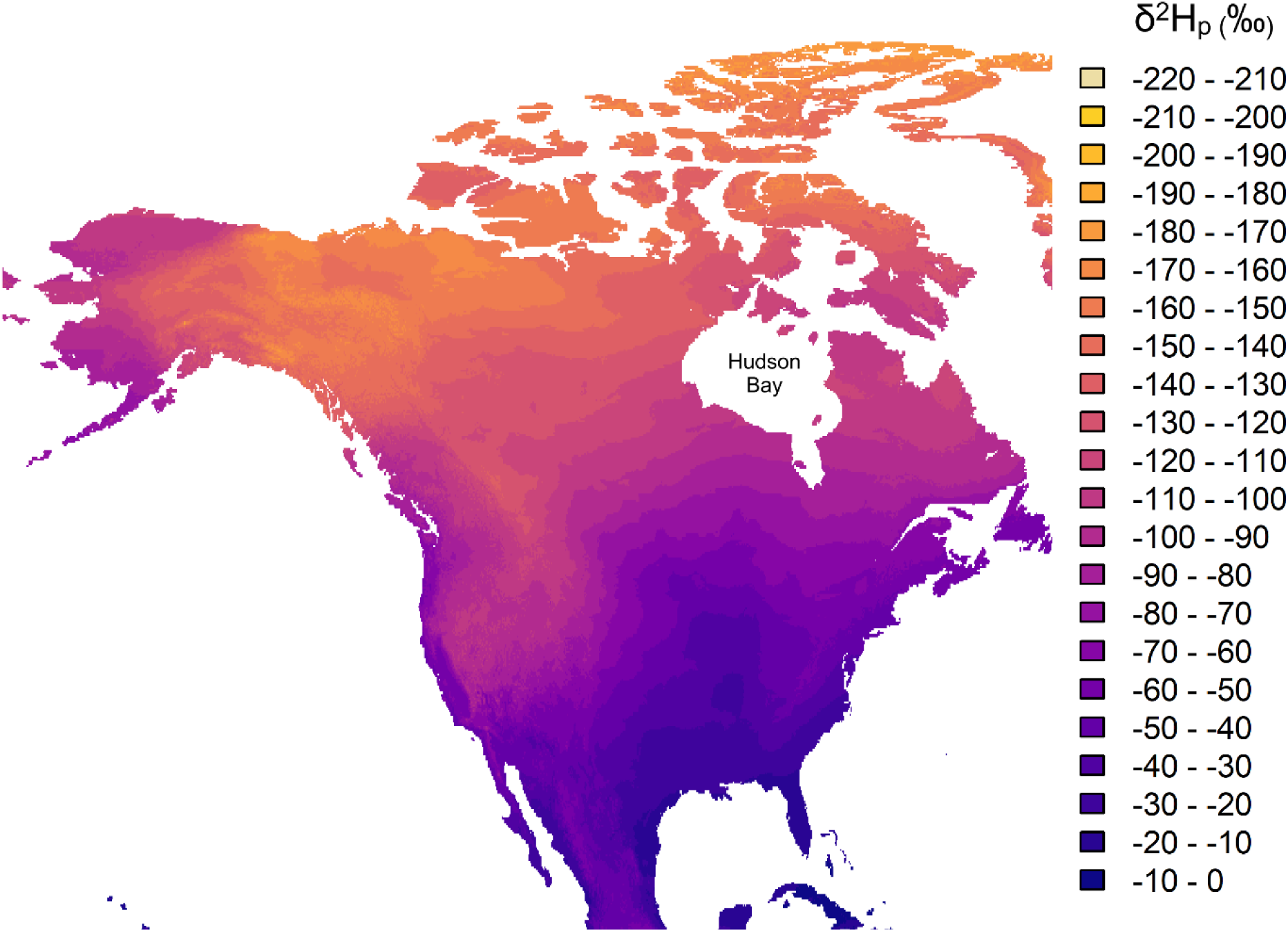
Modelled precipitation deuterium (δ^2^H_p)_ isoscape for North American derived from Bowen et al (2005), depicting mean annual groundwater deuterium levels with 10‰ bandwidths. Note the narrower and non-uniform bandwidths around Hudson Bay in the central and eastern Arctic.

To produce probability surfaces, we estimated the feather tissue measurement error to be 2.0% standard deviation. We used a mask to constrain the posterior probability estimates to fall within the known range of semipalmated sandpiper, extracted from the eBird status and trends products at a 27km^2^ resolution (Fink et al. 2024). We further removed the Alaskan portion of the species’ range from consideration because the underlying precipitation isoscape mirrors values in eastern Canada, adding unnecessary noise to our probability estimates. We assume, based on previous tracking and bill length studies, that birds from the Alaskan breeding populations are not staging in the Canadian Maritimes (Gratto-Trevor et al. 2012, Brown et al. 2017).

For each individual probability surface, we used a 3:1 odds ratio to identify regions with a high probability of origin, meaning that cells with probabilities ≥ 75% were assigned a value of 1 (probable) and all other cells (< 75%) considered improbable (0). Previous studies used either a 2:1 (e.g., Van Wilgenburg and Hobson 2011, Tonra et al. 2015) or 3:1 odds ratio approach (Rushing et al. 2014, Hobson and Kardynal 2023). We chose the less conservative probability threshold for this study to prioritize specificity (fewer false positives) over sensitivity (fewer false negatives) to better highlight regions with the greatest probability of natal origin. However, given our focus on comparing different methods, which odds ratio was used is inconsequential so long as it is consistent across approaches. The produced binary surfaces were summed across individuals to get an overall population origin estimate for all semipalmated sandpipers captured during this study.

### Comparing informative vs flat priors

To compare informative versus flat priors, we used a known-origin reference sample consisting of all passerines breeding in North America, not including aerial insectivores, for the calibration equation to produce the feather isoscape (Table 1). While semipalmated sandpipers are not passerines, this reference choice was made to compare the effect of including informative versus flat priors because it has the greatest geographic coverage and captures the largest among-sample variance in feather deuterium values. Alternative reference sources were evaluated subsequently (see *Testing alternative calibrations*). We fit two competing models maintaining the same general modeling structure as described above. The first included only the range mask and no other prior probability information to constrain posterior probability estimates. The second model included a hierarchal approach, using informative priors derived from citizen science data (eBird; Sullivan et al. 2009) to constrain the probability of origin estimates based on a coarser range mask and a finer-scale breeding occupancy probability surface. Specifically, we used eBird relative breeding occupancy values at a 3km^2^ resolution to weight posterior probability estimates. In other words, for a given region that would be treated as equivalent based on isotopic values alone, the estimated probability of origin will be weighted towards areas with greater predicted breeding occupancy. Since eBird breeding occupancy predictions are model-based, using 79 environmental predictors and accounting for sampling effort (Fink et al. 2020), this approach is effectively applying priors that are based on breeding habitat preferences.

**Table 1.** Datasets of known-origin samples used to build alternative calibration equations to convert the groundwater isoscape (δ^2^H_p)_ derived from average annual precipitation to a feather isoscape (δ^2^H_f)_. Each calibration equation used a combination of different published datasets (see below), or in the case of the Arctic-breeding shorebirds, are samples of known origin from this study. The R^2^ value represents the strength of the association between δ^2^H_p a_nd δ^2^H_f._ See Appendix Table S1 for a full list of species.

| Correction | Species | Samples | Sites | $R^2$ | Citations |
| --- | --- | --- | --- | --- | --- |
| Passerines | 41 | 1209 | 717 | 0.74 | 1, 2, 3, 4, 5 |
| Ground-foraging insectivores | 12 | 256 | 126 | 0.81 | 1, 2, 3 |
| Raptors | 12 | 260 | 254 | 0.65 | 6 |
| Arctic-breeding shorebirds | 4 | 15 | 3 | 0.51 | This study |

Citations: (1) Hobson et al (2012), (2) Hobson and Wassenaar (1997), (3) Hobson and Koehler (2015), (4) Magozzi et al (2021), (5) Chabot et al (2012), (6) Lott and Smith 2006

The probability of origin surfaces produced by both models were compared visually by standardizing posterior probability values to demonstrate differences in areas highlighted as the most likely natal origin for the population. Additionally, we estimated pseudo-precision of the model output, given that we did not have any known-origin samples of semipalmated sandpipers to calculate true model precision or accuracy. For each individual posterior probability density surface, we created a binary surface using a 3:1 odds ratio. We then calculated the proportion of probable origin cells by dividing the number of cell values equal to 1 by the total number of cells (Reese et al. 2019) and subtracting from 1. A higher proportion is equivalent to an assumed greater precision, indicating the model predicted fewer cells with high likelihood of natal origin.

### Testing alternative calibrations

We compared three alternative reference sources to calibrate the feather isoscape. These sources included: 1) all passerines (without aerial insectivores), 2) ground-foraging insectivores, and 3) raptors (Table 1). The passerine reference source was chosen because it has the greatest geographic coverage and therefore captures a greater extent of the variance in environmental deuterium levels. Ground-foraging insectivores were chosen because, in the absence of actual shorebird samples of known origin for North America, their trophic level matches most closely to Arctic-breeding shorebirds. Finally, despite being at a higher trophic level, raptors were included because the reference source also covers a large geographic range, and importantly, regions at higher latitudes than the other sources.

An individual model was fit using a calibration equation derived from each of the three alternative reference sources and breeding origin probability surfaces were estimated using the hierarchical prior structure detailed above (i.e., range constraint and breeding occupancy weighting). We again used a 3:1 odds ratio approach to threshold the prior probability surfaces and summed across individuals within each approach to generate a population-level estimate of natal origin. These surfaces were compared visually and quantitatively. First, each probability surface was subtracted from the other to create a surface where each cell represents the difference in the total number of individuals estimated to have originated from that area, highlighting regions of disagreement between methods. Second, we estimated the weighted centroid for each of the three approaches, where the centroid was weighted by the number of individuals predicted to have originated from each cell. We then calculated pairwise distances among all centroids to represent the extent of spatial differences among core natal origin estimates. Third, we calculated the pairwise Root Mean Squared Estimate (RMSE), where larger relative values indicate greater overall disagreement. We also estimated the coefficient of variation (CV) for each individual cell among the three approaches and mapped these values across the breeding range to indicate regions with the highest among-method variance. Finally, we used the Jaccard Similarity index for each pairwise comparison where 0 indicates complete disagreement, and 1 indicates complete agreement in method outputs.

### Known-origin Arctic-breeding shorebirds

Given the small sample size and relative lack of geographic variation, especially longitudinal variation, we could not compare a calibration using known-origin samples from Arctic-breeding shorebirds alone to the calibration reference sources described above. Instead, we combined our known-origin samples with the passerine reference source in the calibration equation to assess alignment and compared the posterior probability surface of likely origin to that of the passerine method only. This approach essentially evaluates the added value of including Arctic samples of known origin. We used similar quantitative metrics as described above, including RMSE and the Jaccard Similarity index. In addition, we calculated the Euclidian distance of the predicted probabilities between each paired cell to highlight finer scale regions of disagreement.

### Comparison with known geographic relationships

A well-established clinal association exists between bill length and longitude in semipalmated sandpipers once controlling for sex, where individuals breeding in the eastern Arctic have longer bills than those farther west (Gratto-Trevor et al. 2012). Using the probability surfaces produced above, combined with individual measurements of bill length and sex, we tested whether our estimates of breeding origin aligned with published expectations. Specifically, we used breeding origin estimates produced with the eBird prior method and the passerine calibration reference supplemented with known-origin Arctic-breeding shorebird samples. We created a correlational distance matrix among individual probability surfaces and then fit a hierarchical clustering algorithm using a between-group average linkage (UPGMA) method with 1000 bootstrap iterations. This approach results in each individual being assigned to one of a limited number of defined regions, which maximized similarity of probability distributions within clusters and differences among clusters. We summarized bill length within each identified cluster and compared it to published clines. We also fit two variants of a linear model to test for a relationship between cluster and bill length. Bill length was the response variable with cluster ID as the explanatory variable, but one model included an interaction between cluster and sex, while the other was additive. We report significance at the α = 0.05 level.

Statistical analyses were performed using R statistical software (version 4.3.1; R Core Team 2023). Breeding origin assignment models were fit using the ‘assignR’ package (Chao et al. 2020), eBird data was accessed using ‘ebirdst’ (Strimas-Mackey et al. 2025), geospatial analysis used ‘sf’ (Pebesma and Bivand 2023), and hierarchical clustering used ‘isocat’ (Campbell et al. 2020). Data were plotted using the package ‘ggplot2’ (Wickham 2016).

## Results and Discussion

### Informative versus non-informative priors

Using relative breeding occupancy derived from eBird as a prior probability surface constrained estimates of breeding origin and emphasized different regions than a non-informative prior approach (Figure 2). Specifically, using informative priors highlighted Southampton Island and the north-northwest shores of Hudson Bay into the central Arctic. Using non-informative priors also highlighted these regions, but more diffusely, as well as regions farther south than are likely to have dense breeding populations (Figure 2). An estimate of precision based on the relative number of highly likely predicted breeding origin sites indicated greater precision using informative (mean ± SD; 0.797 ± 0.023) versus non-informative priors (0.750 ± 0.060). This translates into an average of 8,989 versus 11,069 cells identified as highly likely (using a 3:1 binary threshold) for the informative prior and non-informative prior method, respectively, or an 18.8% increase in precision. Among-individual variation in total number of predicted origin cells was also over two times greater for the non-informative prior method (SD = 2,642) compared to the informative prior method (SD = 1,006).

**Figure 2.**
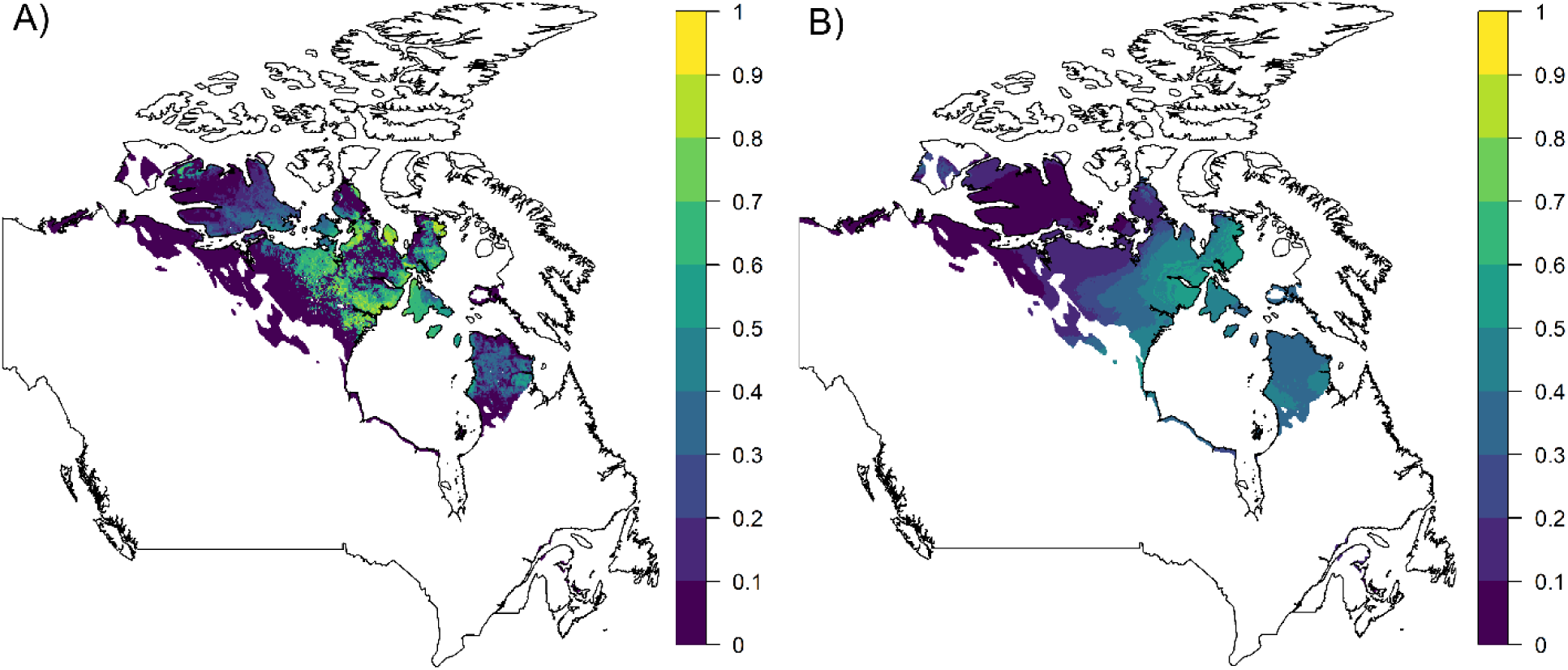
Combined predictions of estimated origin for all 143 semipalmated sandpiper using (A) an occupancy probability raster as a prior and (B) no prior. Estimated probabilities of origin were converted to a binary likely (1) or unlikely (0) using a 3:1 ratio threshold and then summed across individuals. The colour ramp indicates the relative number of individual birds estimated to have originated from that location, standardized between 0 and 1 to equalize comparison. The shaded area is the expected breeding range in Canada derived from eBird at a 27 km^2^ resolution.

Despite being limited in our ability to assess precision and accuracy, our results demonstrate that informative priors can improve origin estimates, at least by focusing on habitat preferences within heterogenous landscapes. The non-informative prior approach produced a more diffuse probability surface, with moderate-to-low probability of population origin (< 50%) across the breeding range. In other words, there was a relatively equal probability of the staging population coming from broad areas representing much of the eastern and central Arctic. Breeding origin predictions at this coarse scale likely underestimate the strength of migratory connectivity and have limited utility for conservation. Using informative priors highlighted distinct ‘hotspots’ within broad isotopic bands, driven by our current understanding of breeding habitat preferences. Here, we are assuming that the underlying relative breeding occupancy probability surface used as a prior is accurate, but it does raise the question of introduced bias from incomplete Arctic breeding data (Reed et al. 2018). While propagated uncertainty is clearly a concern, we argue that the eBird relative occupancy probability estimates are reliable in this context. First, the Arctic PRISM survey, incorporated into eBird, has now achieved complete coverage of the Canadian Arctic (Smith et al. 2025). Second, the eBird data layers are produced using spatiotemporal exploratory models (STEM) that combine observational data and remote-sensing habitat variables to predict occupancy while accounting for spatial biases in coverage (Johnston et al. 2015). This decouples the reliance on abundance counts and prioritizes breeding habitat preferences, in effect down-weighting habitat types that semipalmated sandpiper are unlikely to breed in. Previous research using observational or survey data found that relative abundance estimates can improve origin estimate precision (Hallworth et al. 2013, Rushing et al. 2017); although in some cases reduce accuracy (Reese et al. 2019). Known-origin samples for semipalmated sandpiper collected across their breeding range are required to explicitly quantify accuracy and improve a modeling approach that incorporates priors through an iterative process.

An additional consideration is our focus on hatch-year birds. We chose to sample juvenile birds because we could be sure that their feathers were grown at the breeding site and therefore would be representative of the isotope ratios from the breeding environment. However, if productivity is poorly correlated with occupancy or abundance, assessing only juvenile birds may introduce bias that does not reflect the overall migrating population at the staging site (Hobson et al. 2009). In other words, estimated breeding origin for juveniles may not match the origin of adults using the same staging habitat. While this is not a concern for evaluating different methods, as done here, it must be considered when using prior probabilities based on occupancy to evaluate migratory connectivity. Semipalmated sandpipers conduct a definitive pre-basic molt of head, dorsal and breast feathers on their breeding ground (Gratto 1983, Hicklin and Gratto-Trevor 2020). Therefore, future research could compare predictions from models fit to adults and juveniles separately to better understand differences between age categories and potentially infer spatial variation in productivity across Arctic-breeding populations.

### Alternative isoscape calibrations

We compared predictions across three feather isoscapes derived from reference databases of known-origin samples, representing passerines, ground-foraging insectivores, and raptors (Table 1). Differences between the centroids of predictions were greatest for the calibration equation derived from the raptor data: 1) passerines – raptors (mean, 95% CI; 443 km, 416 – 471 km), and 2) insectivores – raptors (435 km, 402 – 469 km). There was minimal difference between the passerine and insectivore calibration method centroids (59 km, 54 – 63 km). Similarly, RMSE values were greatest for the passerine vs raptor approach (0.850) and insectivore vs raptor approach (0.842), relative to the passerine vs insectivore approach (0.224). Visual comparisons supported these differences, with finer-scale and weaker differences between passerines and insectivores (Figure 3A), compared to using the raptor dataset, which tended to predict breeding origin farther west on average (Figure 3B). The median coefficient of variation across cells among all three approaches was 0.457 (0.159 – 0.861; 10% - 90% quantile), with the greatest disparity tending to occur along the periphery and the extreme east and west (Figure 3C). Finally, the Jaccard Similarity index, ranging from 0 (completely dissimilar) to 1 (identical) was 0.351 (Passerine-Insectivore), 0.214 (Insectivore-Raptor), and 0.110 (Passerine-Raptor), highlighting low similarity overall, but nearly complete disagreement with the raptor approach.

**Figure 3.**
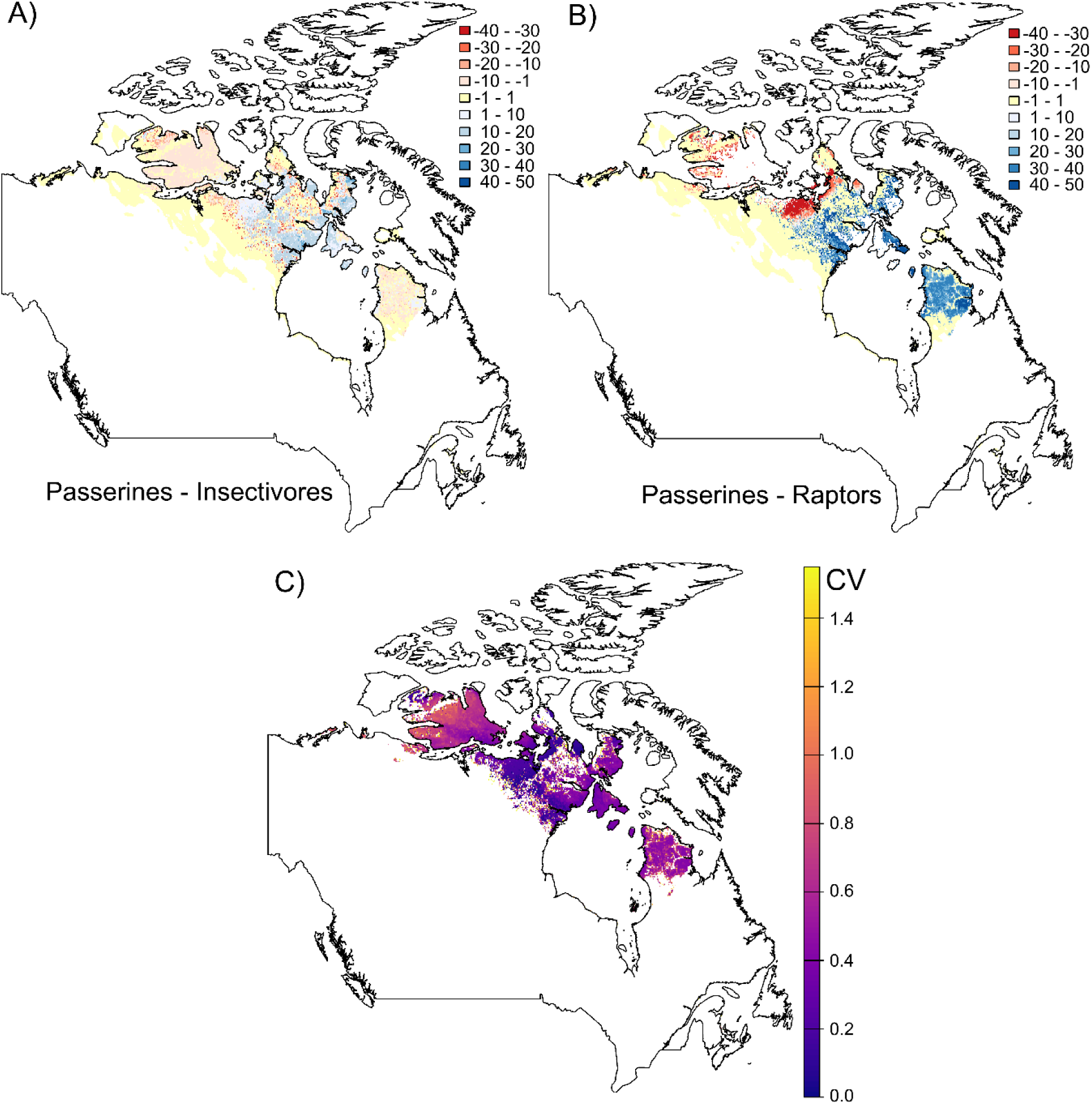
Visual comparison of differences in predicted breeding origin of semipalmated sandpiper, with (A) depicting the difference between the passerine and insectivore calibration and (B) the passerine and raptor calibration. The summed binary estimated origin layers for each method were subtracted from each other to highlight areas of disagreement. Values represent the total difference in predicted individuals with a high probability of originating from the area, with negative values indicating the passerine approach predicted less and positive values meaning the passerine approach predicted more. Panel C depicts the coefficient of variation (CV) among the three approaches, with lighter colours indicating greater disagreement. In all panels, white indicates no predicted occupancy, while in the top two panels yellow generally indicates little-to-no probability of origin (i.e., zero difference between approaches).

As expected, the raptor calibration approach produced significantly divergent estimates of origin. Tissue deuterium levels from species at high trophic levels tend to average out the variance introduced by the effects of individual factors on isotopic values, potentially making them more stable indicators of environmental deuterium levels for a given location (Bump et al. 2007, Hobson et al. 2012). However, isotope ratios still undergo fractionation when moving across trophic levels (Hobson 2011) which would need to be accounted for to produce a calibration equation relevant for Arctic-breeding shorebirds. This requires an extra step, without which, clearly resulted in an erroneous shift in predicted breeding origin farther west.

More interestingly, most comparative metrics indicated only slight differences between using the broader passerine database versus the more restrictive ground-foraging insectivore database to produce a calibration equation. This was expected given that the ground-foraging insectivore database is a subset of the larger passerine database (Table S1). In contrast, the Jaccard Similarity index suggested relatively low agreement between the passerine and ground-foraging insectivore calibration equation approaches (0.351), largely because it is based on exact matches in probability values rather than relative agreement; unlikely in a probabilistic framework. Most distinct disagreements tended to occur at the range peripheries, where we would already expect greater uncertainty based on breeding occupancy probability. Specifically, the ground-foraging insectivore dataset predicted higher probability of origin across a larger extent of the Arctic, both farther west and east, while the passerine approach concentrated in the central-east Arctic. While species included in the ground-foraging insectivore dataset might be a closer trophic level match to Arctic-breeding shorebirds, the geographic variation in known-origin samples is limited compared to the larger passerine dataset (Figure S1). This greater standing variance may allow for more reliable and precise predictions because the Bayesian framework explicitly models uncertainty around the calibration equation line to refine estimates (Wunder 2010, Hobson et al. 2012). In the absence of known-origin samples to properly compare accuracy among alternative calibrations, our comparison indicates prediction stability or low sensitivity to the exact species used to build a calibration equation when estimating origin. Instead, we highlight that the geographic and isotopic variance among known-origin samples used to build the calibration equation are likely more critical than minor trophic level differences.

### Known-origin samples for Arctic-breeding shorebirds

Due to the small sample size and limited spatial variation, using known-origin Arctic-breeding shorebird samples alone to produce a feather isoscape was not possible and represented a relatively poor correlation between the precipitation isoscape (δ^2^H_p)_ and feather deuterium levels (δ^2^H_f_; Table 1). However, the measured feather samples fit relatively well along the best-fit line representing the three primary calibration equations (Figure 4). Using the passerines approach as the default, we compared the relative contribution of adding the known-origin Arctic-breeding shorebird samples to the passerine dataset. The difference between the passerine only and passerine plus Arctic-breeding shorebirds approach was minimal (RMSE = 0.008; Jaccard Similarity index = 0.981). Yet, spatially, Euclidian distances between the two approaches across predicted cells highlighted the greatest disagreement centered around northern Quebec (Figure 5). This suggests that the inclusion of Arctic-breeding shorebird samples shifted predicted breeding origin slightly farther east on average (Figure S2), though predictions were nearly identical generally.

**Figure 4.**
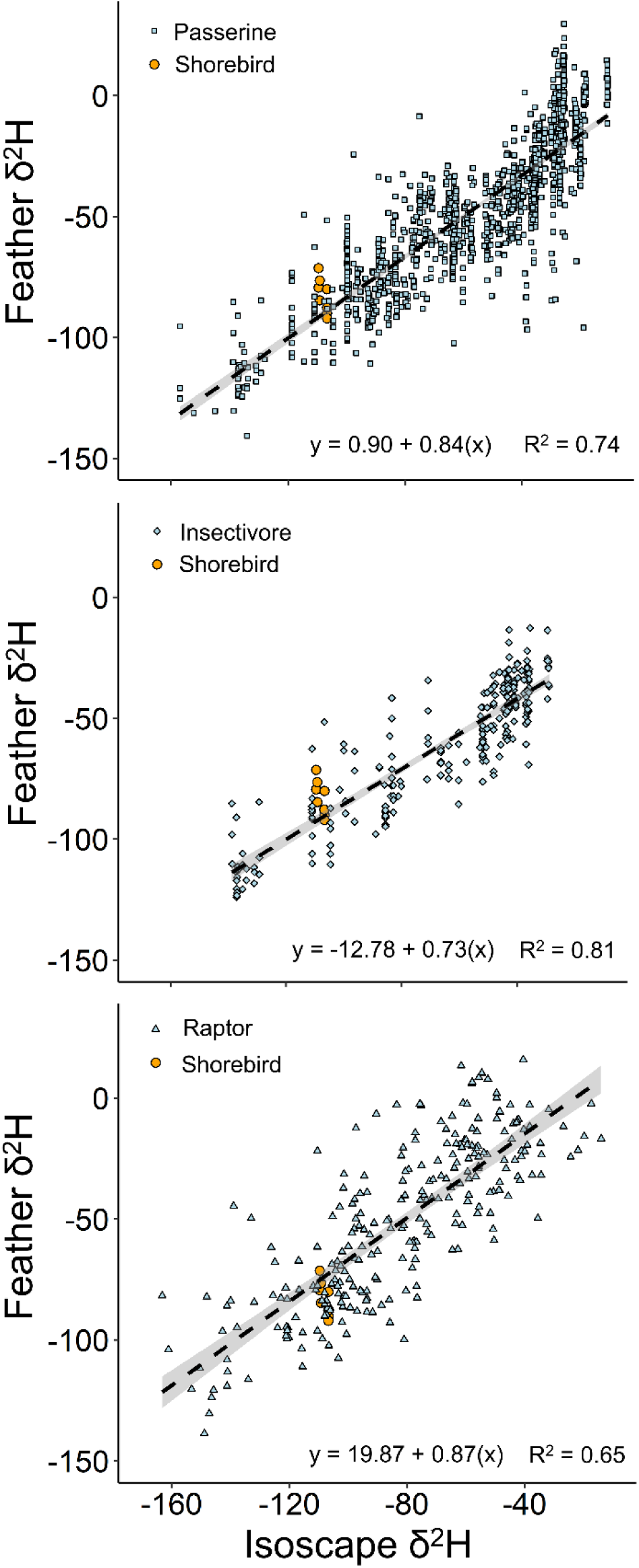
Correlations between groundwater δ^2^H levels and δ^2^H levels from feathers of known origin for passerines, ground-foraging insectivores, and raptors (top to bottom). The orange points depict where our sampled Arctic-breeding shorebird values exist along the best fit line. The linear calibration equation and proportion of variance explained (R^2^) are included in each panel. See Table 1 for citations of the underlying data.

**Figure 5.**
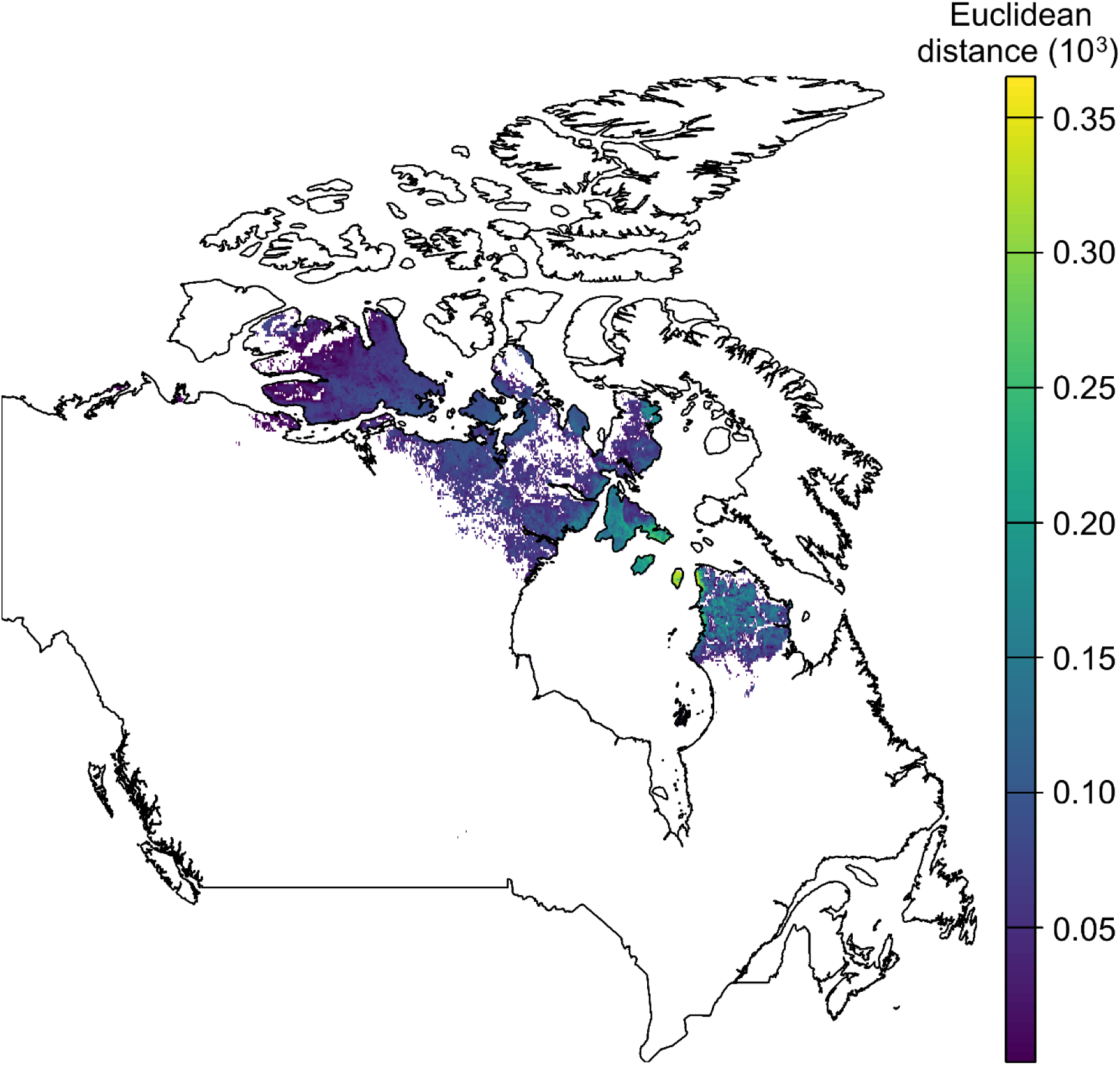
Euclidian distance between estimated breeding origin estimates using the passerine approach versus the passerine plus Arctic-breeding shorebird approach. Lighter colours indicate a greater distance or disparity between approaches at that particular point.

Despite the small sample size of known-origin Arctic-breeding shorebird samples, our results provide some promising takeaways for future work. First, the Arctic feather samples fall along the calibration equation line, particularly for the passerine reference dataset. This strengthens the argument that a calibration equation derived from the overall passerine dataset may produce a sufficiently representative feather isoscape for Arctic-breeding shorebirds. Second, the minor deviation east when incorporating Arctic-breeding shorebird samples indicates that current calibration approaches may be slightly over-estimating the western extent of origin assignments. This should not be an issue for relative comparisons among populations, but it does underline the need for more known-origin samples of Arctic shorebirds to confirm and refine the estimation process. Future work should aim to increase the number of Arctic samples and prioritize longitudinal variance to be more representative.

### Validation with bill length clines

Our hierarchical clustering approach defined two well-separated clusters (Figure 6). Males assigned to breeding origins in the east had longer bill lengths (cluster 1; mean ± SD; 19.3 ± 1.2 mm) relative to the central Arctic (cluster 2; 18.7 ± 0.8 mm), and females demonstrated an even more distinct difference (east: 21.1 ± 1.67 mm, central: 20.1 ± 0.9). Once controlling for sex, the difference was approximately 0.92 mm or 4.7% between clusters (β = -0.69, P < 0.001). Females also had bill lengths that were ∼ 10% longer than males across both clusters (β = -1.52, P < 0.001), aligning with previous research that found regional differences between sexes of 6 – 11% (average = 10%; Gratto-Trevor et al. 2012). There was no evidence for an interaction between cluster ID and sex (F_1,138 =_ 0.11, P = 0.74), indicating that the relationship between clusters was similar for both sexes.

**Figure 6.**
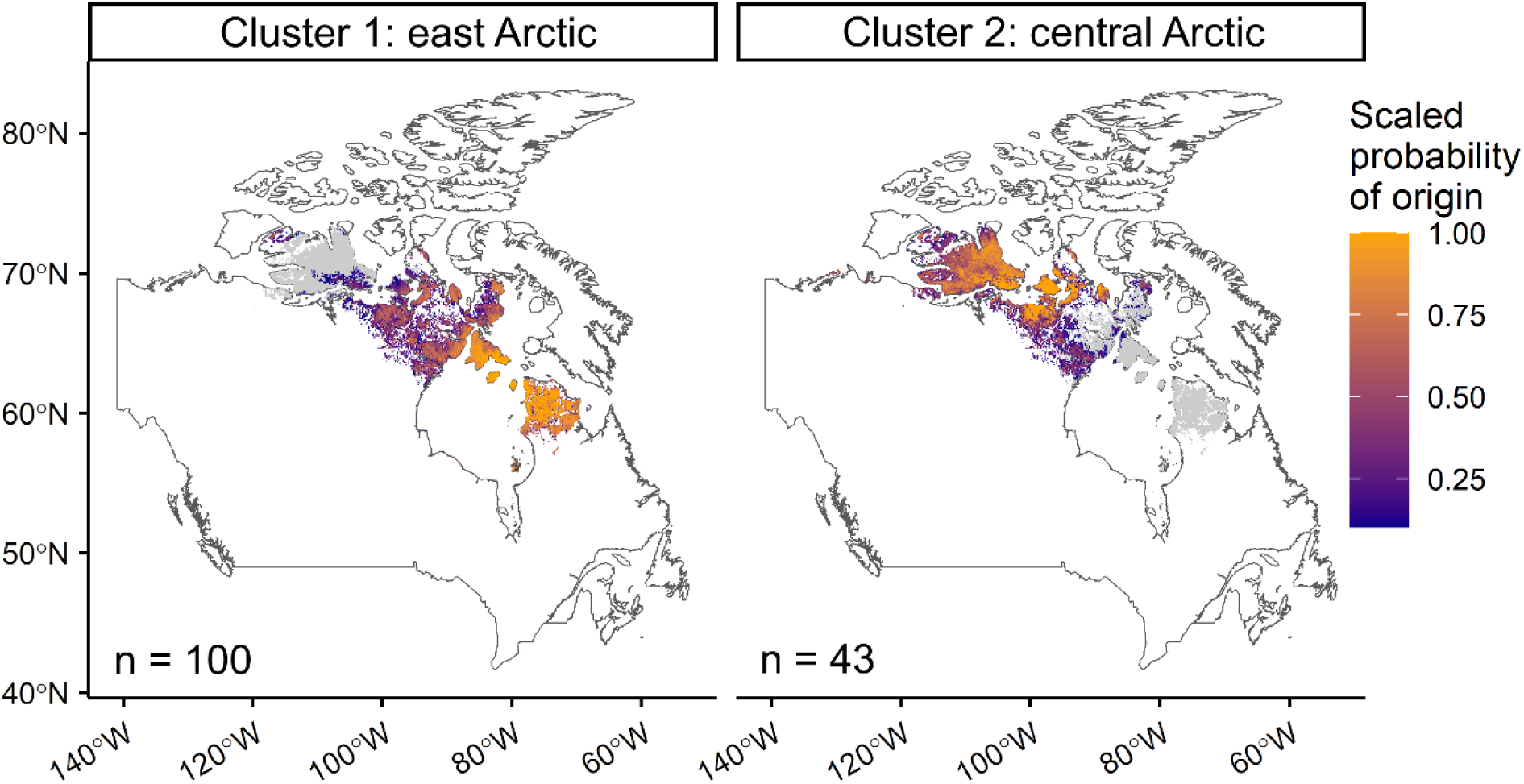
Probability of breeding origin estimates scaled across individuals for the two primary regions identified using a hierarchical clustering algorithm. The clusters represent individuals more likely to have originated from the east Arctic (n = 100) versus farther west in the central Arctic (n = 43). Individuals assigned to each cluster were summarized by sex to assess differences in bill length.

The hierarchical cluster approach differentiating individual by bill length into two broad regions is a relatively coarse scale method of validation. However, this result aligns with the spatial scale that we can assess, given that previous research has binned the origin-bill length relationship into three broad categories (eastern, central, and western Arctic; Gratto-Trevor et al. 2012). Since we only addressed the east and central Arctic in this study by removing Alaska (west Arctic) from consideration, we therefore would only expect two clusters based on bill length. While this method of ground-truthing is not comparable to finer-scale estimates of accuracy from known-origin samples, it does confirm the general expected pattern and that using existing calibration equations in conjunction with occupancy probability priors is sufficient to differentiate among broad regions.

## Conclusion

Given differential population declines among populations of Arctic-breeding shorebirds, being able to accurately assign breeding origin to shorebirds captured at southern staging or non-breeding grounds is of vital importance to evaluating population levels, changes over time, and developing conservation policies. Using stable isotopes to estimate breeding origin for birds captured away from the breeding grounds is a powerful, cost-effective, and scalable approach, particularly for species like Arctic-breeding shorebirds that breed across an expansive and difficult to access region. This study clearly highlights the need for known-origin feather samples from the Arctic and the development of a shorebird-specific calibration equation to better refine origin assignments, echoing previous calls to diversify taxa (Hobson et al. 2014, Reed et al. 2018, Hobson and Kardynal 2023). We strongly recommend building on our preliminary sample of known-origin samples, focusing on increased breadth across shorebird species and longitudinal variation within the Arctic. In the current absence of a shorebird-specific calibration equation, we highlight the utility of a geographically broad reference database at a similar trophic level, like the passerine database used here. We also demonstrate the potential for using eBird occupancy probability surfaces as priors to inform the modeling process based on habitat preferences. We encourage wider adoption of this approach and further research into the potential biases introduced by different prior probability data sources. Combined, the calibration equation and modeling structure we assessed appears to produce reasonable estimates of breeding origin for Arctic-breeding shorebirds and highlights a clear path forward to refining estimates.

## Author contributions

Devin R. de Zwaan and Diana J. Hamilton conceived the idea for the study. Diana J. Hamilton, Rebeca C. Linhart, and Julie Paquet collected field samples. Devin R. de Zwaan conducted the statistical analysis and led writing of the manuscript. All authors contributed to the final version.

## Acknowledgements

We thank Siena Davis, Sara Bellefontaine, Jana Arseneault, Parker Doiron, Veronica Ouellette, Bryn Nurse, Hilary Mann, and other volunteers for assistance with bird capture in Maritime Canada. Mireille Savoie (Mount Allison EASIL lab) ran all deuterium analyses, with assistance from Lindsay Partington and Heidi O’Connor. We thank Jennie Rausch, Paul Woodard, Sarah Neima, Sara Bellefontaine, Doug MacNearney, Andrew Brown, and Anne Ausems for collecting samples from the Arctic and logistical support. Primary funding was provided by NSERC Discovery Grants 2017-05429 and 2023-05990 to DJH, New Brunswick Wildlife Trust Fund, Environment and Climate Change Canada, Career Launcher Internships, Canada Summer Jobs, and Mount Allison University. Arctic work was funded under NSERC Discovery Grant 2019-04017 to EN, as well as ArcticNet, a Network of Centres of Excellence of Canada and the Polar Continental Shelf Program.

## Supplemental Appendix

**Table S1.**
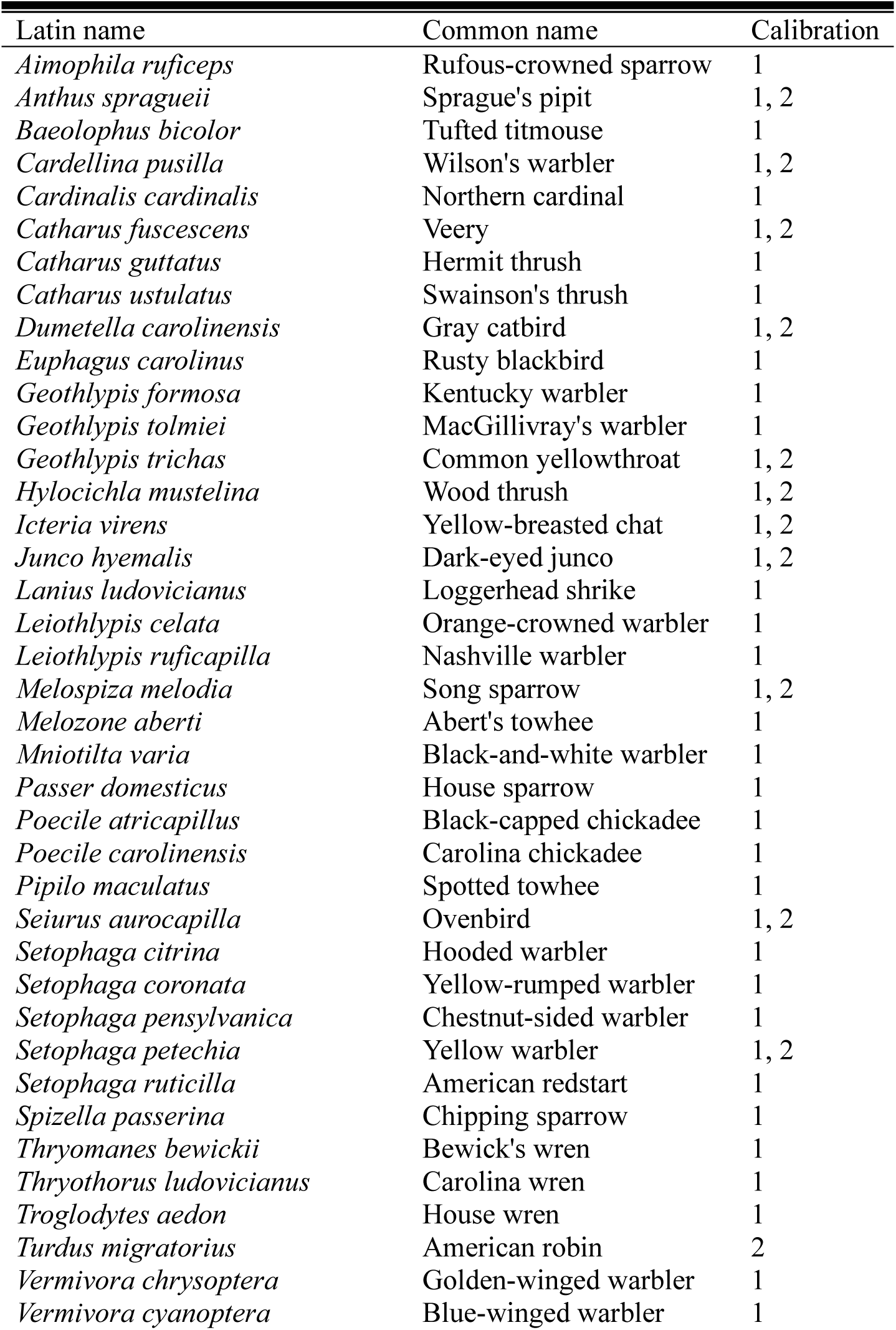
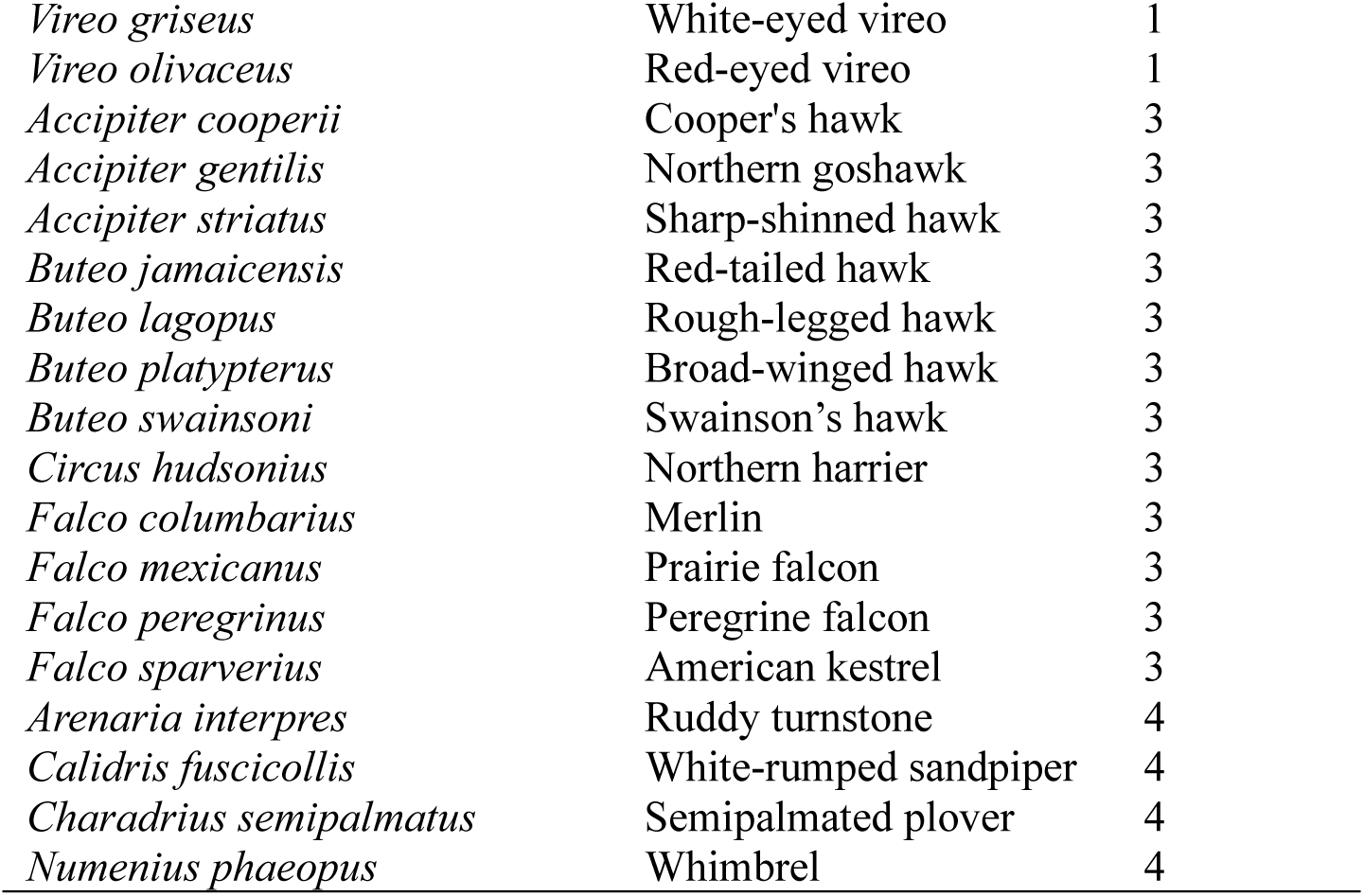
List of species included in each of the four primary datasets with samples of known origin. The calibration dataset includes: (1) Passerines, (2) Ground-foraging insectivores, (3) Raptors, and (4) Arctic-breeding shorebirds.

**Figure S1.**
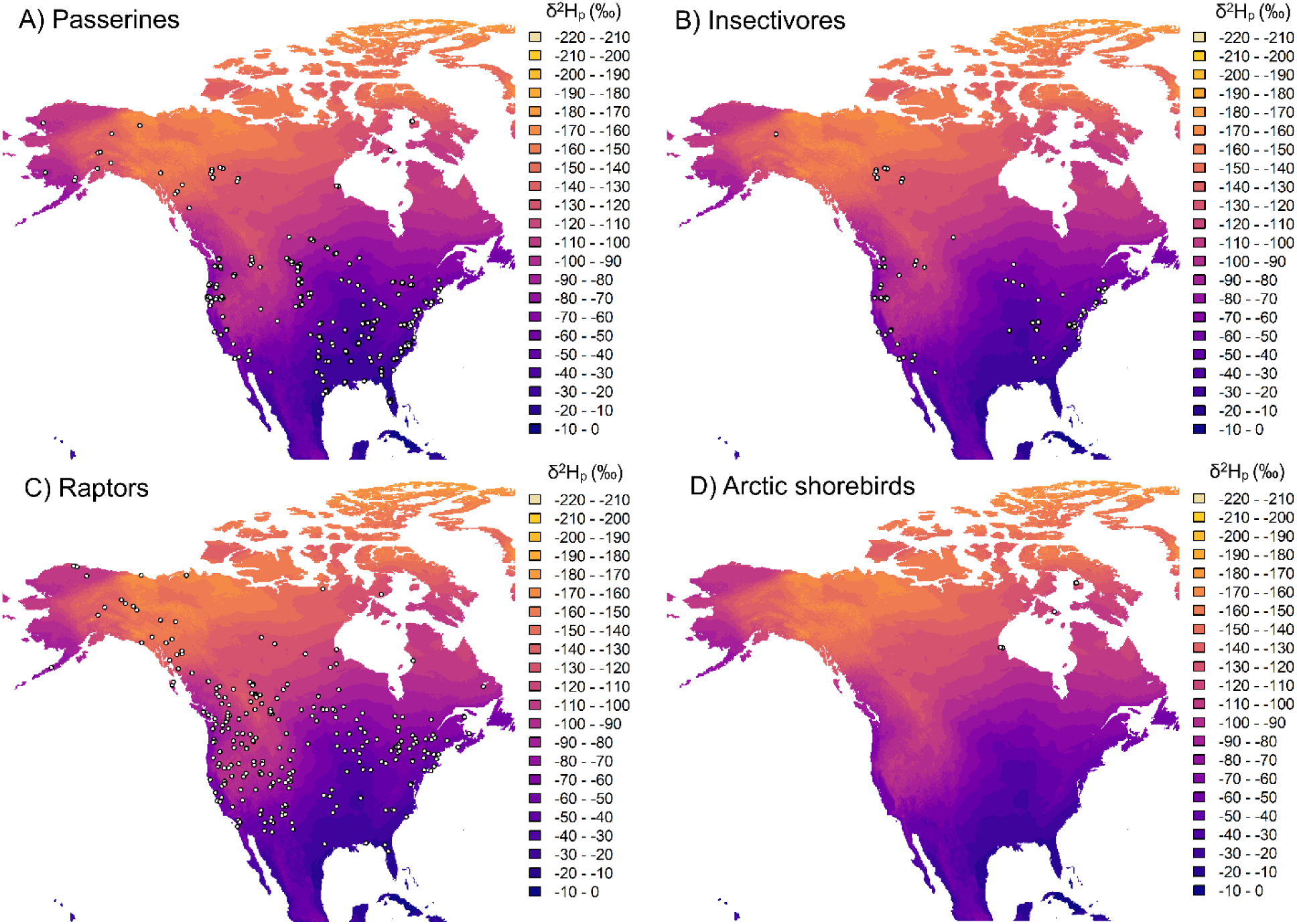
Location of the known-origin samples for each dataset (white dots) overlaid on the modelled precipitation deuterium (δ2Hp) isoscape for North American derived from Bowen et al (2005), depicting mean annual groundwater deuterium levels with 10‰ bandwidths.

